# TREM-1 activation is a key regulator in driving severe pathogenesis of enterovirus 71 infection

**DOI:** 10.1101/682914

**Authors:** Siti Naqiah Amrun, Jeslin J.L. Tan, Natasha Y. Rickett, Jonathan A. Cox, Bernett Lee, Michael J. Griffiths, Tom Solomon, David Perera, Mong How Ooi, Julian A. Hiscox, Lisa F.P. Ng

## Abstract

Hand, foot and mouth disease (HFMD), caused by enterovirus 71 (EV71), presents mild to severe disease, and sometimes fatal neurological and respiratory manifestations. However, reasons for the severe pathogenesis remain undefined. To investigate this, infection and viral kinetics of EV71 isolates from clinical disease (mild, moderate and severe) from Sarawak, Malaysia, were characterized in human rhabdomyosarcoma (RD), neuroblastoma (SH-SY5Y) and peripheral blood mononuclear cells (PBMCs). High resolution transcriptomics was used to decipher EV71-host interactions in PBMCs. Ingenuity analyses revealed similar pathways triggered by all EV71 isolates, although the extent of activation varied. Importantly, several pathways were found to be specific to the severe isolate, including triggering receptor expressed on myeloid cells 1 (TREM-1) signaling. Depletion of TREM-1 in EV71-infected PBMCs with peptide LP17 resulted in decreased levels of pro-inflammatory genes, and reduced viral loads for the moderate and severe isolates. Mechanistically, this is the first report describing the transcriptome profiles during EV71 infections in primary human cells, and the involvement of TREM-1 in the severe disease pathogenesis, thus providing new insights for future treatment targets.

## Introduction

Hand, foot and mouth disease (HFMD) is a febrile illness that predominantly affects infants and young children, and is characterized by rash and blisters on the hands, mouth, feet and bottoms (1–3). Outbreaks of HFMD are caused by human enterovirus group A members (HE-A), mainly coxsackieviruses A16, A6 and A10, and enterovirus 71 (EV71) (1, 4, 5). While often benign and self-limiting (2), the disease can cause cardiopulmonary and neurological complications such as myocarditis, brainstem encephalitis, aseptic meningitis, and neurogenic pulmonary edema, which can be fatal (6, 7). The severe manifestations of HFMD are often associated with cases of EV71 infections, rather than coxsackievirus A16 (8).

EV71 belongs to the *Enterovirus A* species, *Enterovirus* genus, from the *Picornaviridae* family (9, 10). The virus has a positive-sense RNA genome of 7.4 kb and encodes for four structural (VP1-4) and seven non-structural (2A-C and 3A-D) proteins (9). First isolated in California, USA, in 1969 (11), the virus is transmitted via the oral-fecal route, saliva and respiratory secretions (12). Based on the sequence analysis of VP1 gene, the virus can be broadly categorized into six genogroups: A, B, C, D, E and F (7, 10, 13). Whereas genogroup A contains the Californian prototype virus BrCr, genogroups B and C consist of various strains, and are further categorized into sub-genogroups B0-B5 and C1-C5 (10). B and C genogroups are globally distributed, whereas D, and E and F are limited to India and Africa respectively (7, 10). After the almost total eradication of poliovirus, EV71 is now noted as one of the most prevalent neurotropic enteroviruses, and is especially endemic in the Asia-Pacific region (14–17). Currently, the reasons for the emergence of neurological diseases from EV71 infections are still unknown (1), and this poses a significant public health problem as there are no antivirals or vaccines available (18).

In this study, clinical EV71 isolates associated with differing levels of severity (mild, moderate, and severe) collected from an outbreak in 2006 in Sarawak, Malaysia (19), were used to characterize and compare the infection kinetics, virus tropism and immuno-pathogenesis using *in vitro* and *ex vivo* models. In particular, infection kinetics and high-density transcriptomic profiling by RNA-sequencing (RNA-seq) of EV71-infected peripheral blood mononuclear cells (PBMCs) revealed several virus severity-dependent differences. Collectively, this is the first report describing the immune mechanisms involved during EV71 infections in primary human cells, and the identification of novel pathways, such as triggering receptor expressed on myeloid cells 1 (TREM-1) signaling, that are unique and specific to the severe EV71 isolate, and likely contributing to the severe manifestations of the disease.

## Results

### Clinical EV71 isolates are phylogenetically classified based on disease outcomes

Six EV71 isolates were identified from patients during an outbreak of HFMD in Sarawak, Malaysia, in 2006 (20). These patients had differing levels of disease severity, ranging from mild herpangina/HFMD, severe HFMD in the absence or presence of neurological complications, and one fatality (Figure 1A). However, alignment analysis of the different coding sequences of the virus revealed the isolates to be at least 92% identical to one another (Table S1). Moreover, phylogenetic classification of the viruses based on the sequence of the VP1 gene identified them as belonging to the B5 sub-genogroup, and were closely related to strains circulating in Sarawak, Brunei, and Singapore from 2003 to 2008 (Figure S1). Despite sharing 96% VP1 genetic similarity (Table S1), all of the isolates, except isolate 3, were observed to be clustered according to disease severity in the phylogenetic tree (Figure S1).

**Figure 1.**
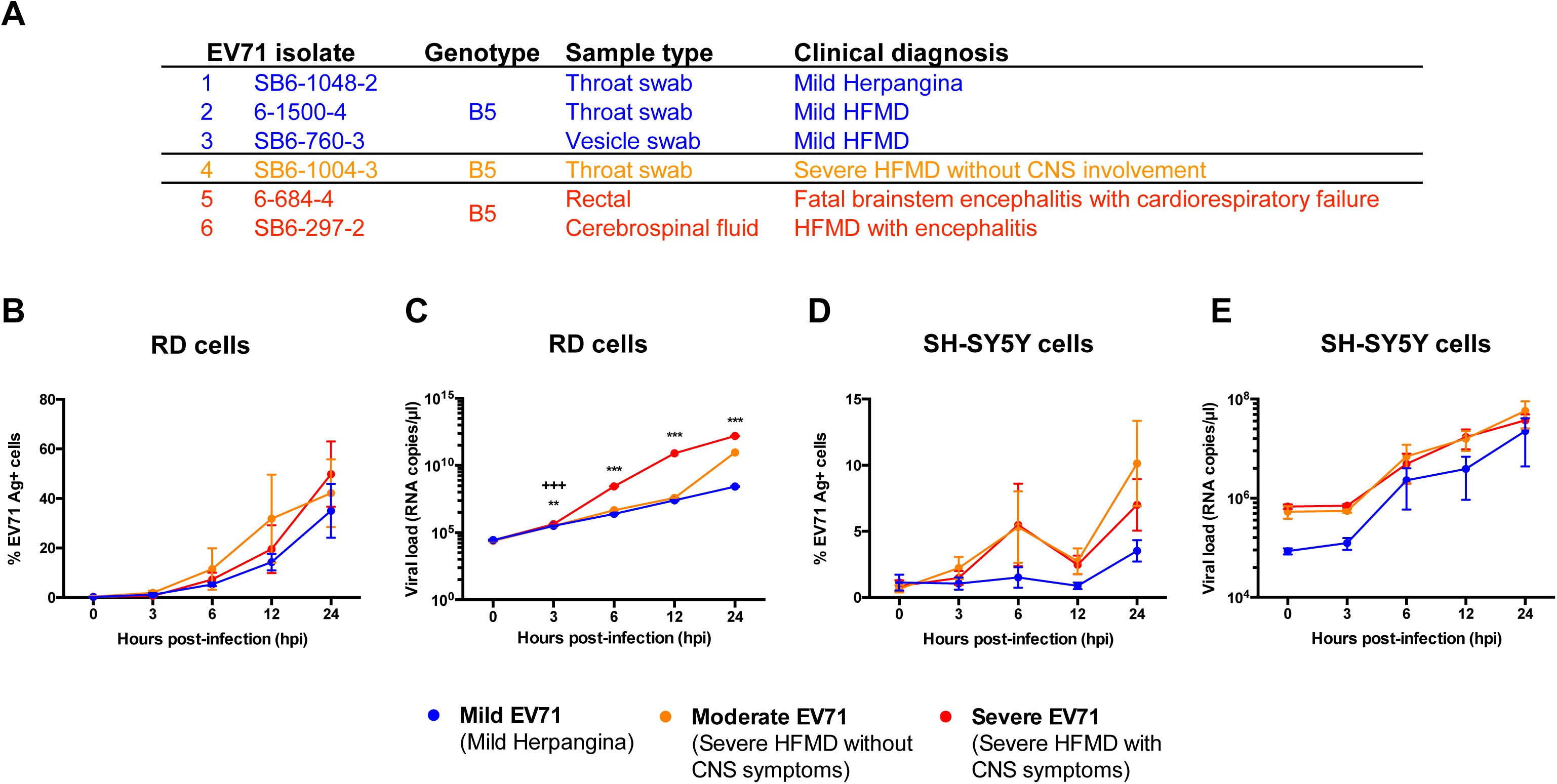
EV71 viral isolates from 2006 outbreak in Sarawak, Malaysia, and its viral kinetics in human rhabdomyosarcoma (RD) and neuroblastoma (SH-SY5Y) cells. (A) Clinical information about the viruses isolated including genotype, symptoms and area virus was isolated from. (B-E) RD and SH-SY5Y cells were infected with mild, moderate and severe isolates of EV71 at MOI 10 and 1 respectively, and harvested at 0, 3, 6, 12 and 24 hours post-infection (hpi). (B-C) Infection kinetics in RD cells via (B) quantification of VP1 antigen by flow cytometry and (C) viral load quantification of VP1 RNA by qRT-PCR. (D-E) Infection kinetics in SH-SY5Y cells via (D) quantification of VP1 antigen by flow cytometry and (E) viral load quantification of VP1 RNA by qRT-PCR. Data are presented as mean ± SEM and representative of 3 independent experiments. Statistical analysis was carried out with Kruskal-Wallis with Dunn’s multiple comparisons test to compare among EV71 isolates at the respective time-points (\*\**p*<0.01; \*\*\**p*<0.001) [Mild vs Severe EV71 (*); Moderate vs Severe EV71 (+)].

### Severe EV71 isolate infects human muscle RD cells to high levels

To investigate if the different EV71 clinical isolates have a different infection profile that could contribute to disease outcome, infection kinetics were compared in human muscle rhabdomyosarcoma (RD) cells and neuroblastoma SH-SY5Y cells at multiplicity of infections (MOI) 10 and 1, respectively. These cell lines have been extensively used in elucidating viral kinetics (7, 21) and pathogenesis of enteroviruses (22, 23). The infection kinetics across a time-course of 0, 3, 6, 12, and 24 hours post-infection (hpi) were determined by measuring infectivity parameters of percentage of intracellular VP1 antigen-positive cells by flow cytometry (Figure S2A) and levels of VP1 RNA load by qRT-PCR. In RD cells, the percentage of infected VP1+ cells among the EV71 isolates 1, 2 and 3 within the mild disease phenotype group were similar by 24 hpi (Figure S2B). Likewise, there were no significant differences between isolates 5 and 6 of the severe phenotype over the course of the time-points (Figure S2C). As such, isolates 1, 4 and 5 were chosen as representatives for further characterization, and were defined as “mild”, “moderate”, and “severe” EV71 isolates respectively.

The severe isolate showed the highest level of infection in RD cells, followed by moderate and mild (Figures 1B and 1C). The percentage of EV71 VP1 antigen in live cells assessed via flow cytometry showed that by 24 hpi, severe EV71-infected RD cells exhibited the highest levels of VP1+ cells, followed closely by moderate EV71, and mild EV71 (Figure 1B). Severe EV71 also showed significantly higher virus replication compared to the mild isolate from 3 to 24 hpi, whereas moderate EV71 had higher VP1 RNA copies than mild EV71 only at 24 hpi (Figure 1C). However, there were no significant differences in percentage of VP1+ cells and viral RNA copies among the three isolates at almost all time-points in SH-SY5Y cells (Figures 1D and 1E).

### Primary human monocytes, B and T cells are the main targets of EV71

Infection kinetics of the different EV71 isolates were next explored in human PBMCs to assess if the associated disease severity affects the ability of the virus to infect the primary cells, which are more biologically relevant than cell lines. In this *ex vivo* model, healthy donor PBMCs were infected with mild, moderate, severe, and heat-inactivated forms of EV71 at MOI 5 over 0, 6, 12 and 24 hpi. PBMCs infected with moderate and severe isolates of EV71 had higher levels of viral VP1 RNA load compared to the mild isolate (Figure 2A). The latter showed lower and limited viral replication throughout the course of infection, whereas moderate EV71-infected PBMCs had the highest viral load in the initial phase, before steadily decreasing over time (Figure 2A). However, PBMCs infected with the severe isolate demonstrated increasing viral load levels till 12 hpi, before dropping off at 24 hpi (Figure 2A). These observed trends were also consistent with the percentage of EV71 VP1 antigen detected from total CD45+ leukocytes (Figures 2B and 2C).

**Figure 2.**
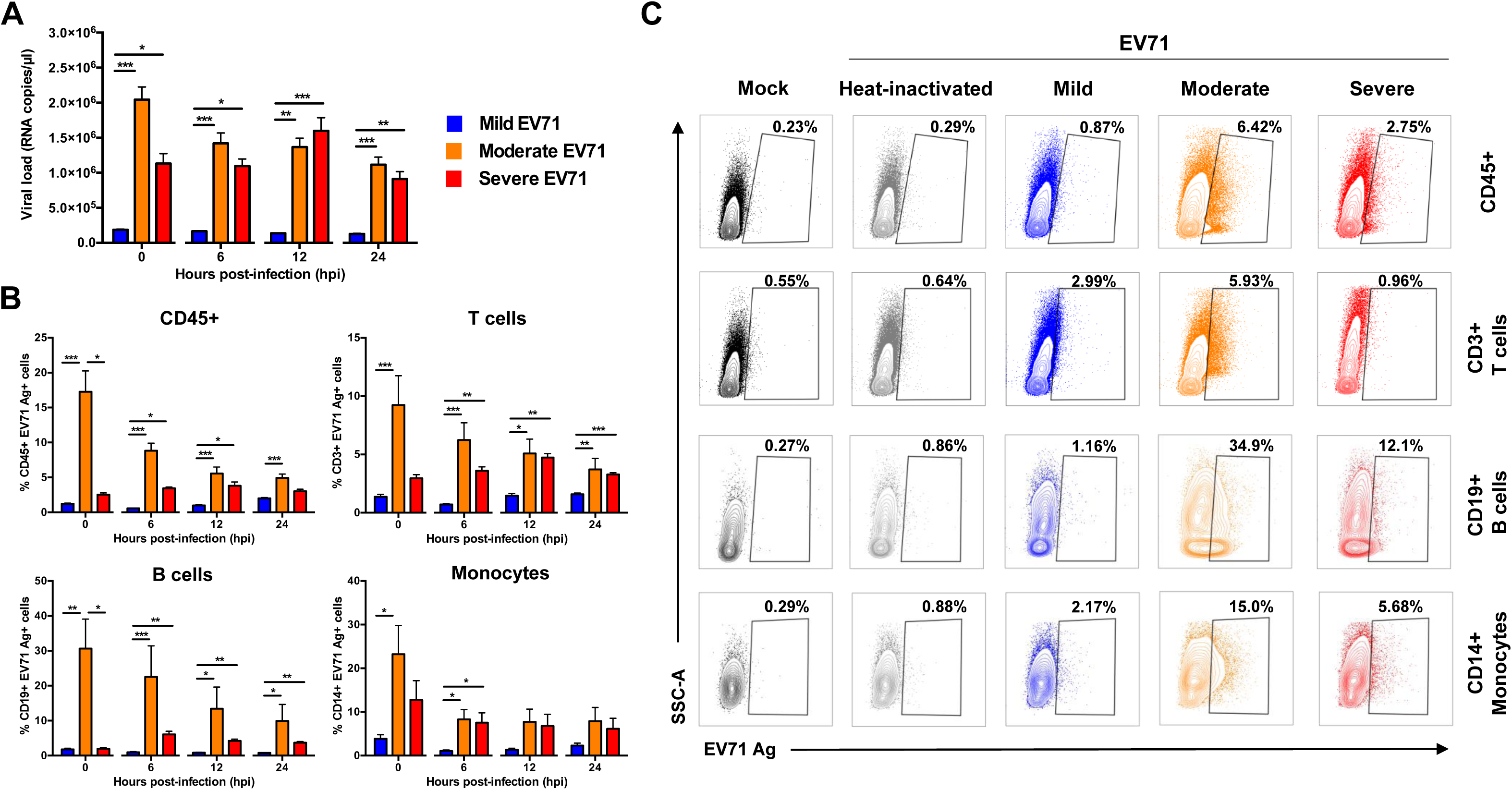
EV71 infection in primary human peripheral blood mononuclear cells (PBMCs). Human primary PBMCs (n=5-9) were infected with mild, moderate, severe and heat-inactivated EV71 isolates at MOI 5, and harvested at 0, 6, 12 and 24 hpi for viral load quantification and flow cytometry. (A) Viral load levels were determined by qRT-PCR. (B-C) Percentage of EV71 Ag+ cells from total CD45+, CD3+ T cells, CD19+ B cells and CD14+ monocytes were determined via flow cytometry. (B) Bar graphs showing the levels of EV71 Ag+ cells from the different cell subsets. Data are presented as mean ± SEM. (C) Illustration of representative contour plots from one donor at 12 hpi. Statistical analysis was carried out with Kruskal-Wallis with Dunn’s multiple comparisons test to compare among EV71 isolates at the respective time-points (\**p*<0.05; \*\**p*<0.01; \*\*\**p*<0.001).

To identify the specific subsets susceptible to EV71 infection, cells were further gated into the different populations: T cells, B cells, monocytes, and natural killer (NK) cells (Figure S2D). T and B lymphocytes, as well as monocytes, were the main susceptible subsets within the PBMCs (Figures 2B and 2C). In contrast, NK cells showed limited and lower levels of infectivity (Figure S2E). Again, PBMCs infected with mild EV71 demonstrated an overall limited percentage of VP1 antigen within the three susceptible subsets (Figure 2B), whereas moderate-infected PBMCs showed highest VP1 percentage in B cells, followed closely by monocytes, and T cells (Figure 2B). On the other hand, PBMCs infected with the severe isolate displayed a slightly different trend, with the highest percentage of VP1 antigen observed in the monocytes, followed by B cells and T cells (Figure 2B). Moreover, there was an increase in the percentage of VP1 antigen in the B and T lymphocytes populations at 6 hpi and 12 hpi respectively, whereas in the monocytes the percentage decreased at 6 hpi but was sustained throughout the course of infection (Figure 2B).

### Transcriptomic profiling reveals a strong induction of host antiviral immune response upon EV71 infection

To identify the possible mediators responsible for the disease severity-dependent differences, RNA-seq was performed on samples of EV71-infected PBMCs from four donors to identify and quantitate mRNA abundance and changes throughout the course of infection at 0, 6, 12 and 24 hpi. Mapped genes of infected conditions were normalized to a heat-inactivated control virus at the respective time-points. The results showed that the number of significant differentially expressed transcripts (false discovery rate [FDR] < 0.05) were different across time-points as well as isolates (Figure 3A). During the course of infection, the greatest change occurred from 0 to 6 hpi for all EV71 isolates, as indicated by the sharp increase in the number of differentially expressed genes (DEGs), and the amount of DEGs continued to show an overall increasing trend from 6 to 24 hpi (Figure 3A). Strikingly, moderate EV71 demonstrated the highest numbers of DEGs throughout infection, followed by severe EV71, and mild EV71 (Figure 3A).

**Figure 3.**
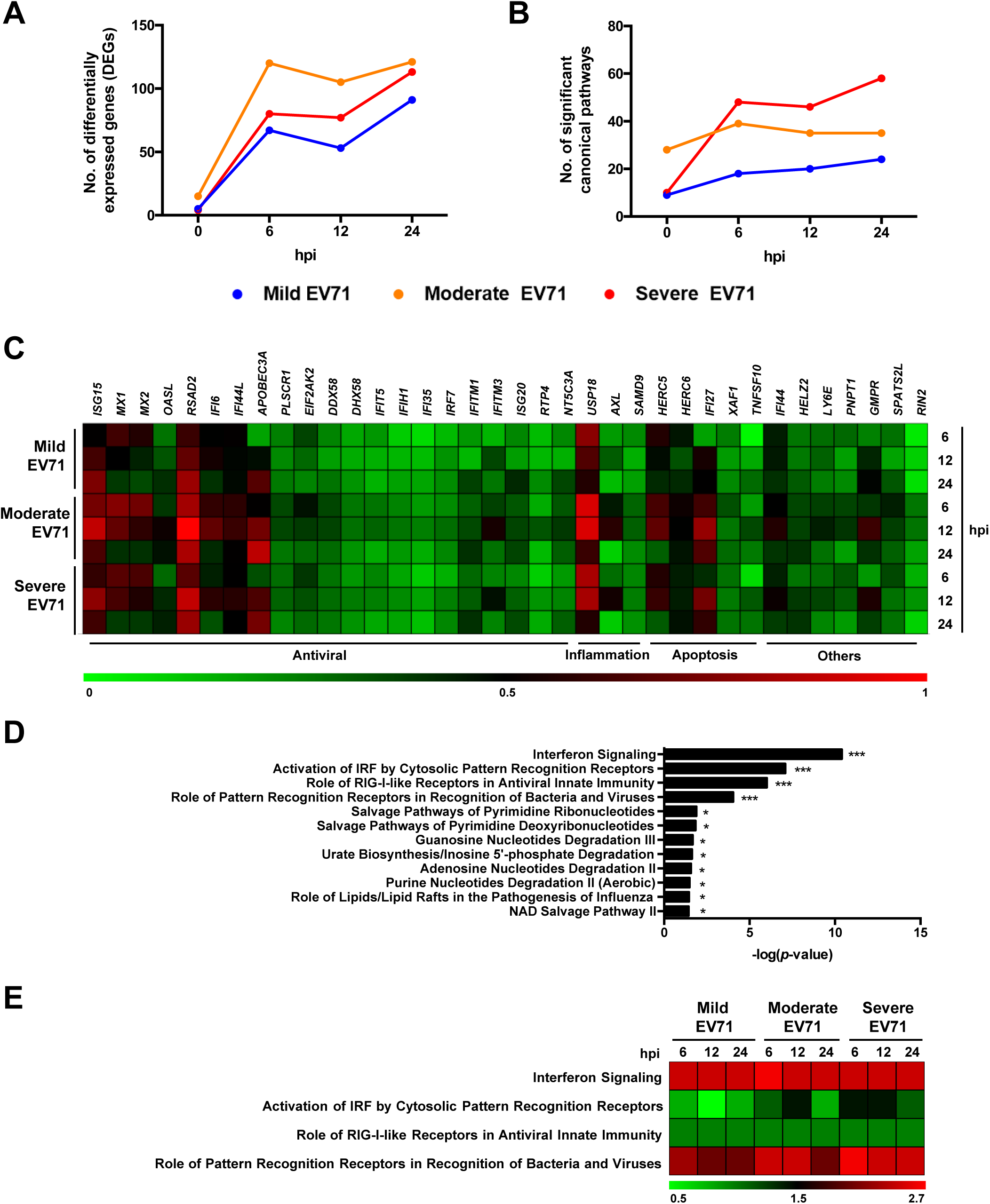
Transcriptomic profiling of human PBMCs during EV71 infection. Primary human PBMCs (n=4) were infected with mild, moderate, severe and heat-inactivated EV71 isolates at MOI 5, and harvested at 0, 6, 12 and 24 hpi for transcriptomic analysis by RNA-seq. Results of infected conditions were compared to heat-inactivated control, and the log_2_-(fold change) values were analyzed in Ingenuity Pathway Analysis (IPA). (A-B) Overall changes of significant (A) differentially expressed genes (DEGs) and (B) canonical pathways identified by IPA over time and in different isolates. (C) Heat-map showing normalized log_2_-(fold change) values of the common DEGs in all infected conditions of 6, 12 and 24 hpi. (D) Significant canonical pathways determined by IPA from the common DEGs in (C) (\**p*<0.05; \*\*\**p*<0.001). (E) Heat-map showing the activation *z*-score values of the top 4 canonical pathways as determined by IPA.

Significant DEGs of the respective isolates and time-points were analyzed in Ingenuity Pathway Analysis (IPA) to identify functional pathways (Figure 3B). The numbers of significantly enriched canonical pathways [−log(*p*-value) > 1.30] were highest for severe EV71-infected PBMCs, especially from 6 to 24 hpi (Figure 3B), despite being presented with fewer differentially expressed transcripts than moderate-infected PBMCs (Figure 3A). PBMCs infected with mild EV71 displayed the lowest number of functional pathways (Figure 3B).

In order to characterize the cellular responses elicited upon EV71 infection, transcripts that were common among the three isolates at time-points 6 to 24 hpi were identified and further analyzed. Regardless of isolate severity, a total of 36 genes were differentially expressed during EV71 infection (Table S2), and these genes are known to be involved in the antiviral interferon response, inflammatory processes, as well as in apoptosis (Figure 3C). Nonetheless, the proportion of fold change for most of the genes differed across time and among the EV71 isolates. Notably, PBMCs infected with moderate EV71 showed a stronger response than the mild and severe isolates, specifically interferon-stimulated gene 15 (*ISG15*), viperin (*RSAD2*), ubiquitin specific peptidase 18 (*USP18*), interferon alpha inducible protein 27 (*IFI27*), and guanosine monophosphate reductase (*GMPR*) (Figure 3C). Next, in order to identify the common functional pathways, the 36 genes were further analyzed in IPA. The results yielded 12 significant canonical pathways (Table S3) and expectedly, the antiviral response and virus recognition formed the top four pathways (Figures 3D and 3E). Based on the activation *z*-scores generated by IPA, the interferon and RIG-I like signaling pathways were equally activated among the EV71 isolates, and across time (Figure 3E). The pathway involving virus recognition by pattern recognition receptors (PRR) was most significantly activated in severe EV71-infected conditions at 6 hpi that persisted till 24 hpi (Figure 3E). However, a different pattern was observed for mild and moderate EV71 infections, in which the activation of this pathway became less pronounced over time (Figure 3E). Similar observations were also seen for the interferon-regulatory factor (IRF) activation pathway (Figure 3E).

### Activation of TREM-1 pathway is specific to severe EV71 infection

Given that the different EV71 isolates belonged to the same sub-genotype B5 (Figure S1), and yet were able to give rise to such varied clinical outcomes in patients (Figure 1A), we then sought to understand this by identifying and comparing the unique DEGs of the EV71 isolates at the respective time-points of 6, 12 and 24 hpi. The three isolates showed distinctive trends (Figure 4A). While the number of unique genes increased slightly from 6 to 13 DEGs (6 to 24 hpi) in the mild-infected condition, the moderate EV71-infected PBMCs displayed an opposite trend, with a decrease in the numbers from 42 to 20 DEGs (Figure 4A). On the other hand, infection of PBMCs with severe EV71 isolate showed an increasing number of DEGs specific to this isolate (4 to 16 DEGs at 6 and 24 hpi) (Figure 4A). Examples of enriched DEGs distinct to each isolate were nuclear protein 1 (*NUPR1*) for mild, chemokine (C-X-C motif) ligand 13 (*CXCL13*) for moderate, and interleukin-6 (*IL-6*) for severe EV71 infections (Table S4). An analysis of the unique DEGs with IPA revealed significant canonical pathways similar to the observations as in Figure 4A (Figure 4B).

**Figure 4.**
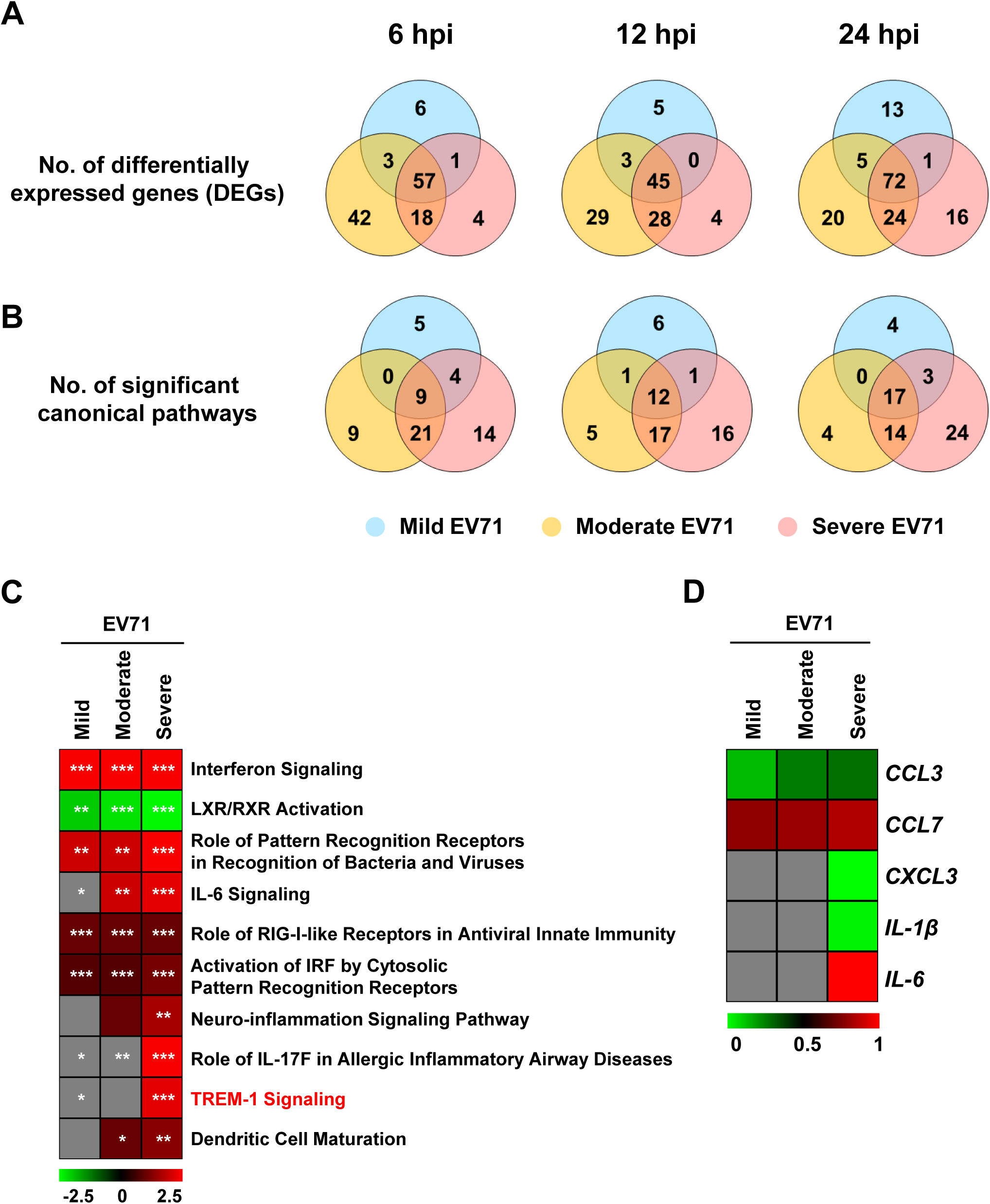
Differences in transcriptomic profiles of human PBMCs infected with different EV71 isolates. Primary human PBMCs (n=4) were infected with mild, moderate, severe and heat-inactivated EV71 isolates at MOI 5, and harvested at 0, 6, 12 and 24 hpi for transcriptomic analysis by RNA-seq. Results of infected conditions were compared to heat-inactivated control, and the log_2_-(fold change) values were analyzed in Ingenuity Pathway Analysis (IPA). (A-B) Venn diagrams illustrating the number of significantly (A) differentially expressed genes (DEGs) and (B) canonical pathways of the different EV71 isolates at 6, 12 and 24 hpi. (C) Heat-map showing the activation *z*-scores of the top 10 canonical pathways at 24 hpi as determined by IPA. Asterisks indicate the significance of the pathway in respective isolates. (D) Heat-map showing normalized log_2_-(fold change) values of the genes involved in TREM-1 signaling pathway at 24 hpi. Grey boxes indicate no values. (\**p*<0.05; \*\**p*<0.01; \*\*\**p*<0.001).

Since the numbers of activated canonical pathways in EV71-infected PBMCs were the highest at 24 hpi (Figures 3B and 4B), the differential genes of each isolate at this particular time-point were input into IPA and comparison analyses were carried out (Table S4). The top 10 canonical pathways, based on the activation *z*-scores as determined by IPA, were selected for further analysis (Figure 4C and Table S5). Remarkably, PBMCs infected with severe EV71 showed a very distinctive response. Pathways such as IL-6, neuro-inflammation, role of IL-17F in allergic inflammatory airway diseases, and TREM-1 signaling processes were significantly and highly activated but were absent or lacking in the mild and moderate EV71-infected conditions (Figure 4C). Notably, the TREM-1 signaling pathway was most distinctive in severe EV71 infection (Figure 4C). The expression of genes involved in this pathway were strongly enriched for *CXCL3*, *IL-1β* and *IL-6* in the severe-infected condition (Figure 4D). Moreover, TREM-1 pathway had not been explored before in the context of EV71 infection, and as such, this pathway was further studied in subsequent experiments.

### Reduced TREM-1 expression mitigates EV71 infection

To verify the role of TREM-1 signaling pathway as an important mediator in EV71 infection, a series of blocking experiments were carried out in which healthy donor PBMCs were pre-treated with 100 ng/ml of synthetic peptide LP17 (Figure S3A). LP17 was previously shown to be able to compete with the ligand and block the TREM-1 receptor, thus inhibiting inflammatory responses (24–26). A scrambled form of the peptide, named sLP17, was used as negative control (Figure S3A). PBMCs were then infected with mild, moderate and severe EV71 isolates at MOI 5, as well as mock-infected, before fresh media containing the final concentration of peptides were added (Figure S3A). In addition, as positive controls, PBMCs were subjected to lipopolysaccharide (LPS) treatment in the presence of peptides. To validate that the blocking was efficient, total RNA samples of peptide- and LPS-treated PBMCs were subjected to gene expression studies by qRT-PCR (Figure S3B). LPS controls with LP17 peptide treatment resulted in decreased expression levels of the genes of interest, especially *IL-6* and *CCL7*, indicating a successful TREM-1 inhibition (Figure S3B).

Similarly, in the EV71-infected conditions, a reduced gene expression profile was observed for the moderate and severe EV71 isolates (Figure 5A). However, the intensity of TREM-1 inhibition was more pronounced in moderate isolate infection when compared to severe-infected condition (Figure 5A). On the other hand, mild EV71 showed minimal changes in the relative level of gene expression, suggesting that TREM-1 activation in this isolate could be too low to detect for any inhibition (Figure 5A). The impact of EV71 viral replication upon TREM-1 inhibition was then assessed by qRT-PCR. Consistent with the minor effects observed in the gene expression results, infection with mild EV71 showed no differences in the number of viral VP1 RNA copies under the blocked and control states (Figure 5B). However, treatment with LP17 to inhibit TREM-1 resulted in a lower viral VP1 RNA load in both the moderate and severe EV71 infections (Figure 5B).

**Figure 5.**
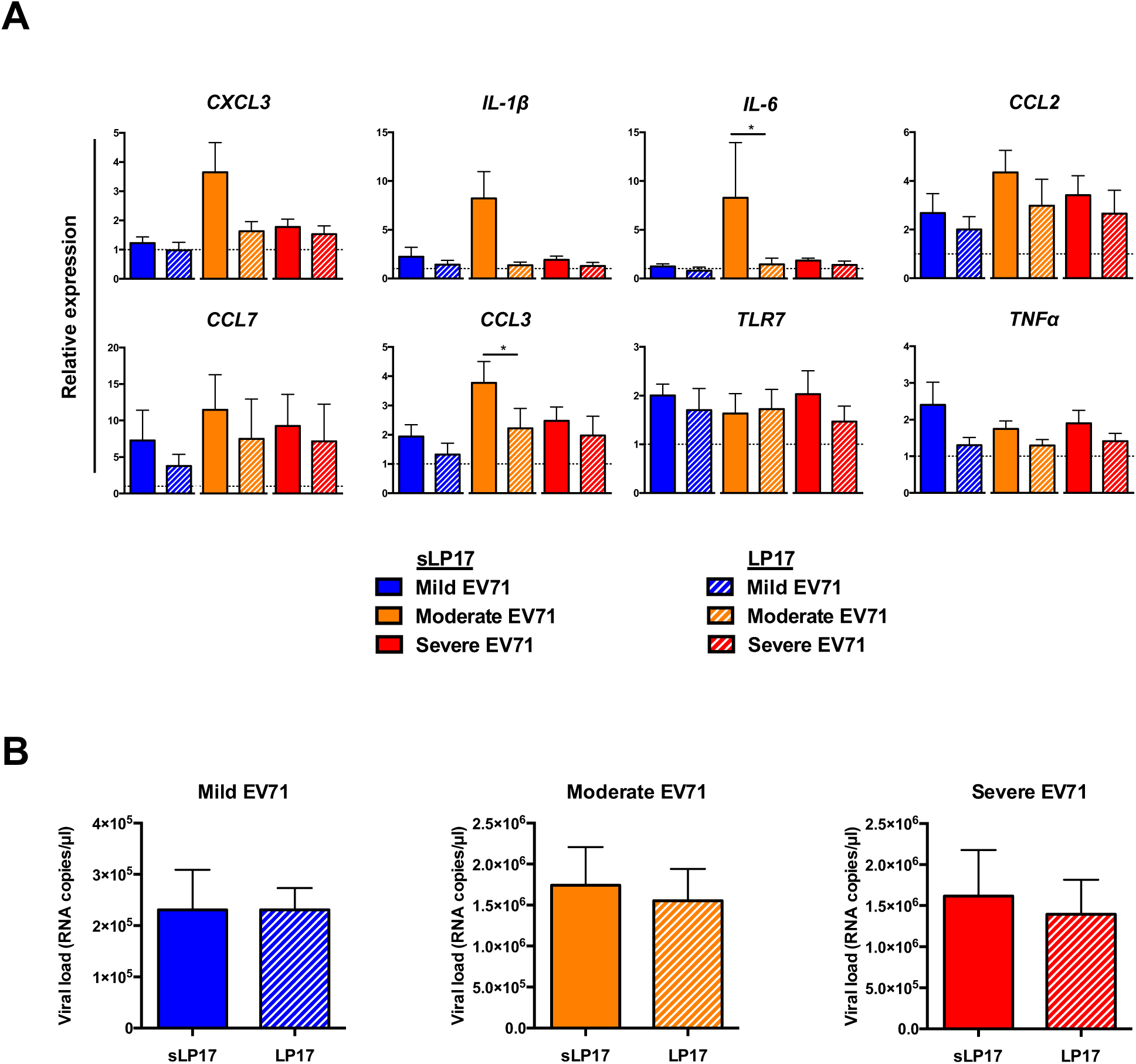
Blocking of TREM-1 by peptide LP17 decreased viral load during EV71 infection. Human primary PBMCs (n=7) were pre-treated with 100 ng/ml of LP17 or the control sLP17 peptide, before infection with mild, moderate, or severe EV71 isolates at MOI 5. Mock-infected PBMCs were used as negative control. Cells were replenished with fresh media containing respective peptides and harvested at 24 hpi for gene expression and viral load quantifications. (A) Total RNA samples were extracted and gene expression studies were performed via qRT-PCR using 10 ng/µl of RNA. Relative expression levels of genes were calculated by comparing the infected conditions to the respective mock. Bar graphs illustrating the expression levels of genes involved in the TREM-1 pathway. Dotted line indicates the baseline levels from mock. (B) Viral load levels were quantified by qRT-PCR of the VP1 RNA copies. Data are presented as mean ± SD. Statistical analysis was carried out with Wilcoxon matched-pairs signed rank test (\**p*<0.05).

## Discussion

Patients suffering from EV71 infections can present different clinical manifestations and severity. Previous studies have suggested that outcomes of EV71 infections could be influenced by the genetic makeup of the virus, as well as the immune status of the patient (19, 27–30). Alignment of the various genes of the isolates revealed that non-structural gene 3A was highly dissimilar between the moderate and mild isolates (92% identity) (Table S1). The 3A protein has been shown to be important for virus RNA replication, with its involvement in the formation of replication organelles (31–33). It was recently reported that recruitment of phosphatidylinositol 4-kinase III*β* (PI4KB) by the interaction of host factor acyl-CoA binding domain containing 3 (ACBD3) and EV71 3A facilitated viral replication (34), thus suggesting that perhaps certain nucleotide differences in the moderate isolate could lead to its increased virulence. Nonetheless, all EV71 isolates were highly genetically identical overall (at least 96%). Yet, viral severity-dependent effects were observed in muscle RD cells and in PBMCs. Moderate and severe EV71 isolates demonstrated high levels of infection and replication, while mild EV71 showed lower infectivity. However in neuronal SH-SY5Y cells, mild and moderate EV71 exhibited similar levels of replication kinetics as the severe isolate, suggesting that the mild and moderate isolates could potentially cause neuro-pathogenesis if the blood-brain barrier permeability is compromised. The results support the importance of the virus’ genetic component in determining virulence and in shaping clinical outcomes in patients.

Intriguingly, between the moderate and severe EV71 infections in PBMCs, the latter exhibited a steady increase in infection and replication over time while the former showed a strong infection only in the initial phase. This was also supported in the transcriptomic results, in which the viral recognition pathway by PRR remained activated specifically for severe EV71, thus emphasizing the ongoing replication of virus over time. In addition, several pathways such as neuro-inflammation and IL-6 signaling were identified to be strongly associated or unique to severe EV71 infection. Cytokines IL-1*β* and IL-6 are the common denominators of these pathways and are therefore likely to play a role in the severe pathogenesis of EV71. In earlier reports, levels of IL-1*β*, IL-6 and TNF*α* were found to be significantly elevated in fatal EV71 patients with encephalitis and pulmonary edema, when compared to patients with no complications (29, 35–37). Production of such inflammatory cytokines are mediated by monocytes, macrophages, as well as T and B lymphocytes (37), and in agreement with this, these immune cells were the subsets shown to be the most susceptible to EV71 in this study and elsewhere (38–40).

The link between pro-inflammatory response and severe EV71 outcome is also further strengthened with the identification of TREM-1 activation, which is unique and distinct to the severe isolate. TREM-1 is a cell surface receptor that is highly expressed on monocytes and neutrophils, and is known as a critical player in the amplification of inflammation via the production of pro-inflammatory mediators (41). TREM-1 can also exist as a soluble variant, with its expression levels indirectly proportional to that of its receptor form (41). To date, TREM-1 modulation in viral infections has been described for dengue, Marburg (MARV) and Ebola (EBOV) viruses among others (26, 42). In dengue-infected patients, the expression of TREM-1 on neutrophils was significantly reduced in the acute phase of the disease, with high levels of soluble TREM-1 in sera (42). On the other hand, MARV and EBOV were able to activate TREM-1 on human neutrophils *in vitro*, and the blocking of TREM-1 receptor by LP17 peptide treatment resulted in diminished production of TNF*α* and IL-1*β* (26). Although a decrease in the expression of pro-inflammatory mediators was also observed in this study, the effect of the treatment was most significant for the moderate isolate of EV71. Moreover, although a lower viral load was observed in the blocked conditions of moderate and severe infections, results were not significant when compared to the control condition. Therefore, inhibition of TREM-1 specifically targeted the downstream immune response after virus infection. Previously, TREM-1-deficient mice infected with either *Leishmania major*, influenza virus or *Legionella pneumophila* were able to demonstrate reduced morbidity and immune-associated pathologies, with minimal effect on pathogen clearance (43). Another plausible explanation could be attributed to the host genetic factors, as donor-to-donor variations were observed in our study. Gene polymorphisms within TREM-1 have been linked to the development of inflammation and sepsis (44), and it has been reported that there is a positive association between patients carrying the *TREM-1* rs2234237T allele and the development of severe malaria, due to the presence of higher levels of soluble TREM-1 in the plasma of these patients (45). Given that polymorphisms in EV71 receptors, scavenger receptor class B member 2 (SCARB2) and P-selectin glycoprotein ligand-1 (PSGL-1), have been shown to affect EV71 susceptibility and infection respectively (46), genomic variations within TREM-1 could also therefore play a role in modulating EV71 infection and outcomes.

In conclusion, this study characterized and compared the cellular responses elicited by EV71 isolates of differing clinical severity and revealed the novel involvement of TREM-1 in the pathogenesis of severe EV71. Importantly, TREM-1 expression levels could serve as a prognostic marker for severe EV71 infections in patients, and might point the way towards future therapies.

## Materials and Methods

### Ethics approval

Human blood apheresis cones were obtained from healthy volunteers with written informed consent, in accordance with the Declaration of Helsinki for Human Research, and with guidelines from the Health Sciences Authority of Singapore (reference number: 2017/2512).

### Cells and viruses

Human rhabdomyosarcoma (RD; ATCC CCL-136) and neuroblastoma (SH-SY5Y; ATCC CRL-2266) cell lines were cultured in Dulbecco’s Modified Eagle Medium (DMEM) and Roswell Park Memorial Institute (RPMI) (HyClone) respectively, supplemented with 10% fetal bovine serum (FBS; Hyclone) at 37°C with 5% CO_2_. EV71 viral isolates used in this study were originally isolated from six patients from the 2006 outbreak in Sarawak, Malaysia (accession numbers MN053430, MN053431, MN053432, MN053433, MN053434 and MN053435). Viruses were propagated for three passages and titered in RD cells, and stored at −80°C. The same passage number of viruses were used for infection experiments. Heat-inactivation of virus was done at 65°C for 45 min.

### Tissue culture infective dose 50 (TCID50) assay

Viral titers were determined in RD cells via TCID50 assay. Briefly, RD cells (1.5 × 10^4^) were seeded in 96-well microtiter plates overnight before infection with serial dilutions of EV71 isolates in serum-free DMEM. After 1.5 h, cells were refreshed with DMEM with 10% FBS and incubated for 4 days at 37°C. Cells were fixed with 10% formalin (Sigma-Aldrich) before being stained with 0.5% crystal violet (Sigma-Aldrich). Titer was calculated using Reed-Muench method (47).

### Genome sequencing and phylogenetic analysis

Viral RNA of the EV71 isolates was extracted using QIAmp Viral RNA Mini Kit (Qiagen) as per manufacturer’s instructions. 30 ng of viral RNA samples were subjected to cDNA synthesis using Illumina TruSeq® RNA sample preparation kit version 2 (Low-Throughput protocol) according to manufacturer’s protocol, except for the following modifications: 1. Skip mRNA purification and start with RNA fragmentation 2. use of 12 PCR cycles, 3. two additional round of Agencourt Ampure XP SPRI beads (Beckman Coulter) to remove >600bp double stranded cDNA. The length distribution of the cDNA libraries was monitored using DNA 1000 kits on the Agilent bioanalyzer. All samples were subjected to an indexed SE sequencing run of 1×51 cycles on an Illumina MiSeq (18 samples/lane). EV71 sub-genogroup B phylogenetic tree was generated by maximum-likelihood analysis of complete VP1 nucleotide sequences aligned using MUSCLE in MEGA. The tree was rooted to the prototype genogroup A strain. Sequences were identified by GenBank accession, year of isolation, and country of origin. Tree robustness was evaluated by bootstrap analysis using 1000 pseudo-replicate sequences. Major clades of bootstrap values of more than 75% were indicated at relevant branch nodes.

### *In vitro* infection of cell lines

RD and SH-SY5Y cells (1 × 10^6^) seeded in 60 mm dish were infected with the EV71 isolates prepared in serum-free media at multiplicity of infection (MOI) 10 and 1, respectively (23, 48–51). After 1.5 h, virus inoculum was removed and replaced with media containing 10% FBS. Cells were incubated at 37°C and harvested at time-points 0, 3, 6, 12, and 24 hours post-infection (hpi). Mock-infected controls were cells without any virus infection.

### *Ex vivo* treatment and infection of human peripheral blood mononuclear cells (PBMCs)

PBMCs were first isolated from a blood apheresis cone by gradient centrifugation using Ficoll-Hypaque method as previously described (52, 53). PBMCs (2 × 10^6^) were then infected with EV71 viral isolates at MOI 5 in serum-free Iscove’s Modified Dulbecco’s Medium (IMDM; Gibco) (50). After 2 h, virus inoculum was removed and replaced with IMDM with 10% human serum (HS; Innovative Research Inc). Cells were incubated at 37°C and harvested at time-points 0, 6, 12, and 24 hpi. Mock-infected controls were cells without any virus infection, whereas heat-inactivated controls were cells infected with heat-inactivated virus. For TREM-1 blocking experiments, isolated PBMCs were first pre-treated with 100 ng/ml of LP17 peptide (LQVTDSGLYRCVIYHPP) or its scrambled form, sLP17 (TDSRCVIGLYHPPLQVY) (Pepscan Systems) as previously described (24, 25), in serum-free IMDM. After 30 min incubation at 37°C, PBMCs were then infected with EV71. Subsequently, peptide-virus solutions were removed and cells were refreshed with 100 ng/ml of sLP17 or LP17 peptides in IMDM and 10% HS. Mock-infected, and LPS-treated PBMCs (50 ng/ml; Sigma-Aldrich) were used as negative and positive controls, respectively.

### Viral RNA extraction and viral RNA quantification

Viral RNA from 140 μl of culture supernatant was extracted using QIAmp Viral RNA Mini Kit (Qiagen) according to manufacturer’s instructions. Viral load quantification was performed with a VP1-targeted qRT-PCR adapted from Tan and colleagues (54), with forward and reverse primers (GAGAGCTCTATAGGAGACAGT and GAGAGCTCTATAGGAGACAGT respectively), and an additional novel lab-designed FAM-labeled molecular beacon probe (ACCCACAGGTCAAAACACACA) taken from consensus sequence of the six EV71 isolates. RT-PCR reactions were prepared using QuantiTect Probe RT-PCR Kit (Qiagen), with final concentrations of primers and probe used were 400 nM and 200 nM respectively. The thermal cycling conditions were: 50°C for 30 min, 95°C for 15 min, followed by a 2-step cycle of a cycle at 94°C for 15 min, and 45 cycles at 55°C for 1 min on 7900HT Fast Real-Time PCR System (Applied Biosystems). Assay exclusivity of EV71-VP1 was confirmed by testing viral RNA extracted from dengue viruses (DENV-1 to DENV-4), Chikungunya virus (CHIKV), and O’nyong’nyong virus (ONNV). Analytical sensitivity was also determined using quantitated EV71 RNA transcripts. The lower limit of detection was estimated as 10 copies for the VP1 gene target. Copy numbers of EV71 RNA were determined using the Ribogreen RNA-specific Quantitation Kit (Invitrogen). RNA transcripts from 10 to 10^9^ copies were performed in pentaplicates to construct a standard curve to estimate EV71 copy number in samples.

### Flow cytometry

RD and SH-SY5Y cells from *in vitro* infections were stained with LIVE/DEAD Fixable Aqua Dead Stain (Life Technologies), fixed with 1X BD FACS Lysing Solution (BD Biosciences), and permeabilized with 1X BD FACS Permeabilizing Solution 2 (BD Biosciences). Subsequently, cells were stained with EV71 VP1 protein specific mouse monoclonal antibody (clone 10F0; Abcam), and counter-stained with fluorophore-tagged goat anti-mouse IgG (H+L) antibody (Invitrogen), before being acquired with BD FACSCanto II (BD Biosciences). PBMCs from *ex vivo* infections were stained and fixed similarly as above, and permeabilized with 0.5% Triton X-100 (Sigma-Aldrich). Cell surface staining was performed with mouse anti-human CD45 (clone HI30; BioLegend), CD3 (clone SP34-2; BD Biosciences), CD19 (clone SJ25C1; eBioscience), CD14 (clone M5E2; BD Pharmingen), CD16 (clone 3G8; BioLegend), CD56 (clone AF12-7H3; Miltenyi Biotec), CD11c (clone 3.9; BioLegend), HLA-DR (clone L243; BioLegend), followed by EV71 VP1 staining and its counter-stain, as above. Cells were acquired with BD LSRFortessa (BD Biosciences). Flow cytometry data were analyzed with FlowJo version 9.3.2 (Tree Star, Inc).

### Total RNA extraction

Total RNA from cells of PBMCs infections were extracted using RNeasy Micro Kit (Qiagen) according to manufacturer’s instructions, and the concentrations quantified with Nanodrop 1000 spectrophotometer (Thermo Scientific).

### Gene expression

Total RNA samples were diluted to a concentration of 10 ng/μl prior to gene expression studies by qRT-PCR using QuantiFast SYBR green RT-PCR kit (Qiagen) in 7900HT Fast Real-Time PCR System (Applied Biosystems) (55, 56) according to manufacturer’s protocol. The C_t_ values for *CXCL3*, *IL-1β, IL-6*, *CCL2*, *CCL7*, *CCL3*, *TLR7*, *TNFα* and *GAPDH* (housekeeping gene) were determined. The fold change relative to mock-infected samples for each gene was determined using the ΔC_t_ method and Microsoft Excel 2016 (55, 56). The sequences of the primers used are in Table S6.

### RNA-seq and differential gene expression analysis

The methods for RNA-seq and differential expression were previously described (57–59) and are briefly described below.

### RNA-seq

RNA samples were DNase-treated using Ambion Turbo DNA-free Kit (Invitrogen), purified using Agencourt Ampure XP beads (Beckman Coulter), followed by a Ribozero treatment using the Epicentre Ribo-Zero Gold Kit (Human/Rat/Mouse) (Ilumina), and purified again with Ampure XP beads. Successful depletion was then quality tested using Qubit 4 Fluorometer (Thermo Fisher) and Agilent 2100 Bioanalyzer (Agilent Technologies), and all of the depleted RNA was used as input material for the ScriptSeq v2 RNA-Seq Library Preparation protocol. Following 14 cycles of amplification, the libraries were purified using Ampure XP beads. Each library was quantified using Qubit, and the size distribution was assessed using the AATI Fragment Analyzer (Advanced Analytical). Final libraries were pooled in equimolar amounts, and the quantity and quality of each pool was assessed by the Fragment Analyzer and subsequently by qPCR using the Illumina Library Quantification Kit (Kapa Biosystems) on a Light Cycler LC480II (Roche) according to manufacturer’s instructions. The template DNA was denatured according to the protocol described in the Illumina cBot User guide and loaded at 12 pM concentration. To improve sequencing quality control 1% PhiX was spiked-in. Sequencing was carried out on three lanes of an Illumina HiSeq 2500 with version 4 chemistry, generating 2 × 125 bp paired-end reads.

### Bioinformatics analysis

Briefly, base calling and de-multiplexing of indexed reads were performed by CASAVA version 1.8.2 (Illumina) to produce 30 samples from the 5 lanes of sequence data, in fastq format. The raw fastq files were trimmed to remove Illumina adapter sequences using Cutadapt version 1.2.1 (60). The option “-O 3” was set, so that the 3’ end of any reads which matched the adapter sequence over at least 3 bp was removed. The reads were further trimmed to remove low quality bases, using Sickle version 1.200 with a minimum window quality score of 20. After trimming, reads shorter than 50 bp were removed. If both reads from a pair passed this filter, each was included in the R1 (forward reads) or R2 (reverse reads) file. If only one of a read pair passed this filter, it was included in the R0 (unpaired reads) file. The reference genome used for alignment was the human reference genome assembly GRCh38. The reference annotation was downloaded from the Ensembl ftp site (ftp://ftp.ensembl.org/pub/release-77/gtf/homo_sapiens/Homo_sapiens.GRCh38.77.gtf.gz). The annotated file contained 63,152 genes. R1/R2 read pairs were mapped to the reference sequence using TopHat2 version 2.1.0 (61) which calls the mapper Bowtie2 version 2.0.10 (62).

### Differential gene expression and functional analysis

Mapped reads were further analyzed using edgeR version 3.3 (63) to calculate normalized counts per million (CPM), to identify genes differentially expressed between infected and heat-inactivated conditions. The log_2_-(fold change) values of such genes were uploaded into Ingenuity Pathway Analysis (IPA; Qiagen) and subsequently used for gene ontology and pathway analysis. IPA was performed to identify canonical pathways, and comparison analysis was used to determine the most significant pathways across the different isolates and time-points. Activation *z*-scores and *p*-values associated with the identified functional pathways were determined by IPA. Heat-maps were generated using TM4 Multi-Experiment Viewer (64).

### Statistical analysis

All statistical analyses were performed using GraphPad Prism 7 (GraphPad Software). For cell lines and PBMCs infection experiments, Kruskal-Wallis with Dunn’s multiple comparison test was used to compare the EV71 isolates at each time-point. For peptide-PBMCs experiments, Wilcoxon matched-pairs signed rank test was used in the comparison of LP17 and control sLP17 peptides in the respective infected condition. *P* values less than 0.05 are considered to be statistically significant.

## Acknowledgements

We thank Fok-Moon Lum, Guillaume Carissimo, Wendy W.L. Lee and Wearn-Xin Yee of Singapore Immunology Network (SIgN); Alicia Tay and Josephine Lum from SIgN Immunogenomics; Ivy Low, Seri Mustafah and Anis Larbi from SIgN Flow Cytometry platforms for their assistance during the study. We are grateful to the donors for their participation in the study.

This study was supported by core research grants provided to SIgN by the Biomedical Research Council (BMRC), Agency for Science, Technology and Research (A*STAR), Singapore. SIgN Immunomonitoring, Immunogenomics and Flow Cytometry platforms are supported by BMRC IAF 311006 grant and BMRC transition funds #H16/99/b0/011. The research was also funded by the National Institute for Health Research Health Protection Research Unit (NIHR HPRU) in Emerging and Zoonotic Infections at the University of Liverpool in partnership with Public Health England (PHE) and Liverpool School of Tropical Medicine (LSTM) to and directly supported NYR, JAH and TS. The views expressed are those of the author(s) and not necessarily those of the NHS, the NIHR, the Department of Health or Public Health England.

## Author contributions

JAH and LFPN conceptualized the study. SNA, JJLT, NYR, JAC, DP, MHO performed the experiments. SNA, JJLT, NYR, JAC, BL, TS, JAH and LFPN analyzed the data. SNA, JJLT, NYR, JAC, BL, DP, MHO, TS, JAH and LFPN wrote the manuscript. All other authors were involved in sample collection, processing and analysis, and/or logistical support, and approved the final version of the manuscript.

