## Supplementary material for "TREM-1 activation is a key regulator in driving severe pathogenesis of enterovirus 71 infection": Figure S1

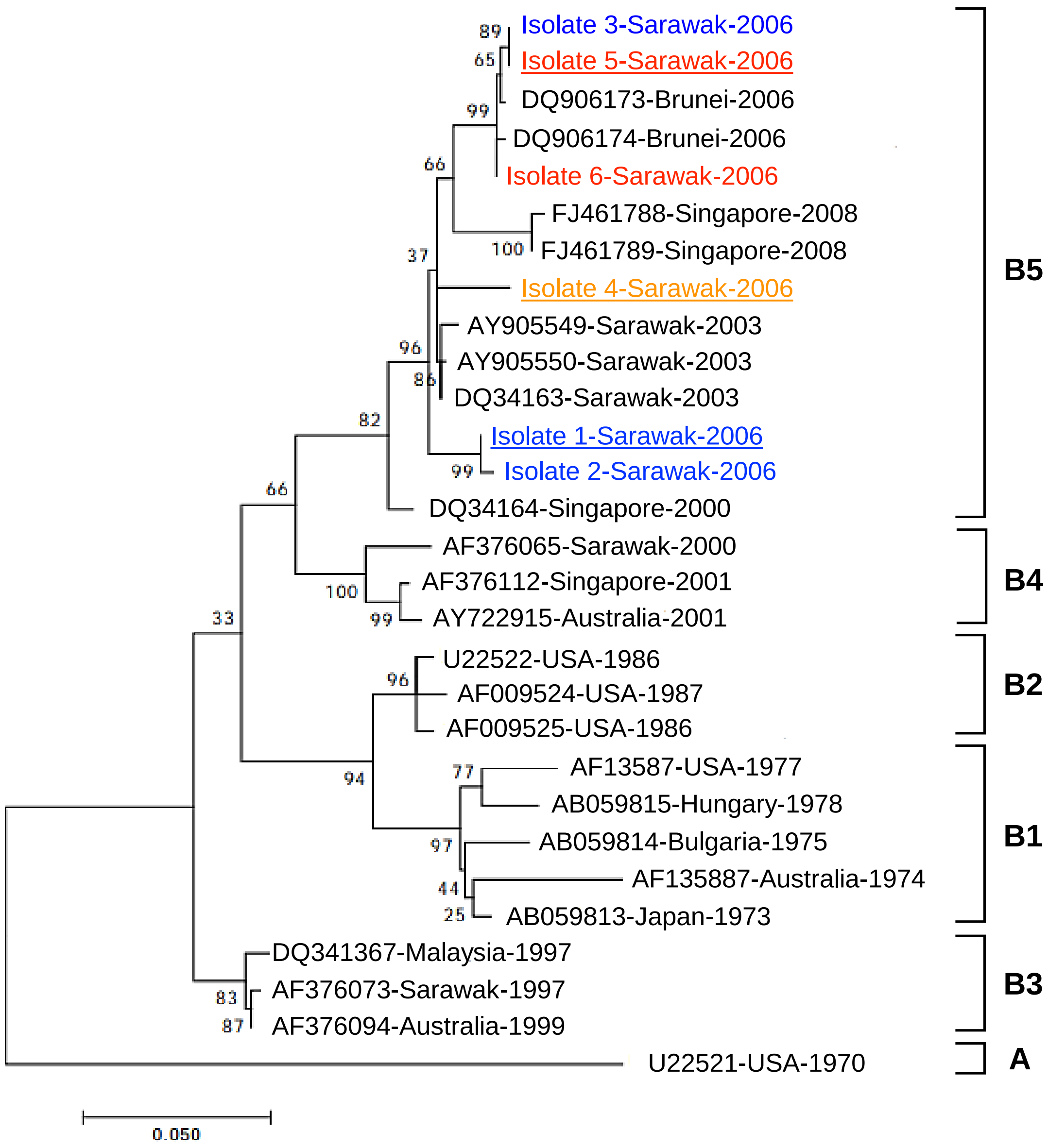


**Figure S1. Phylogenetic classification of EV71 isolates from an outbreak in Sarawak in 2006.** EV71 sub-genogroup B phylogenetic tree generated by maximum-likelihood analysis of complete VP1 nucleotide sequences aligned using MUSCLE in MEGA. The tree was rooted to the prototype genogroup A strain. Sequences are identified by GenBank accession, country of origin and year of isolation. Viruses in this study are colored according to their disease severity, and underlined are viruses that were used for characterization. The robustness of the tree was evaluated by bootstrap analysis using 1000 pseudo-replicate sequences. Bootstrap values >75% of major clades are indicated at relevant branch nodes. All branch lengths are drawn to scale and a measurement of relative phylogenetic distance is provided by the scale at the bottom of the tree.

**Figure S1 Amrun *et al*., 2019**
