## Supplementary material for "TREM-1 activation is a key regulator in driving severe pathogenesis of enterovirus 71 infection": Figure S2

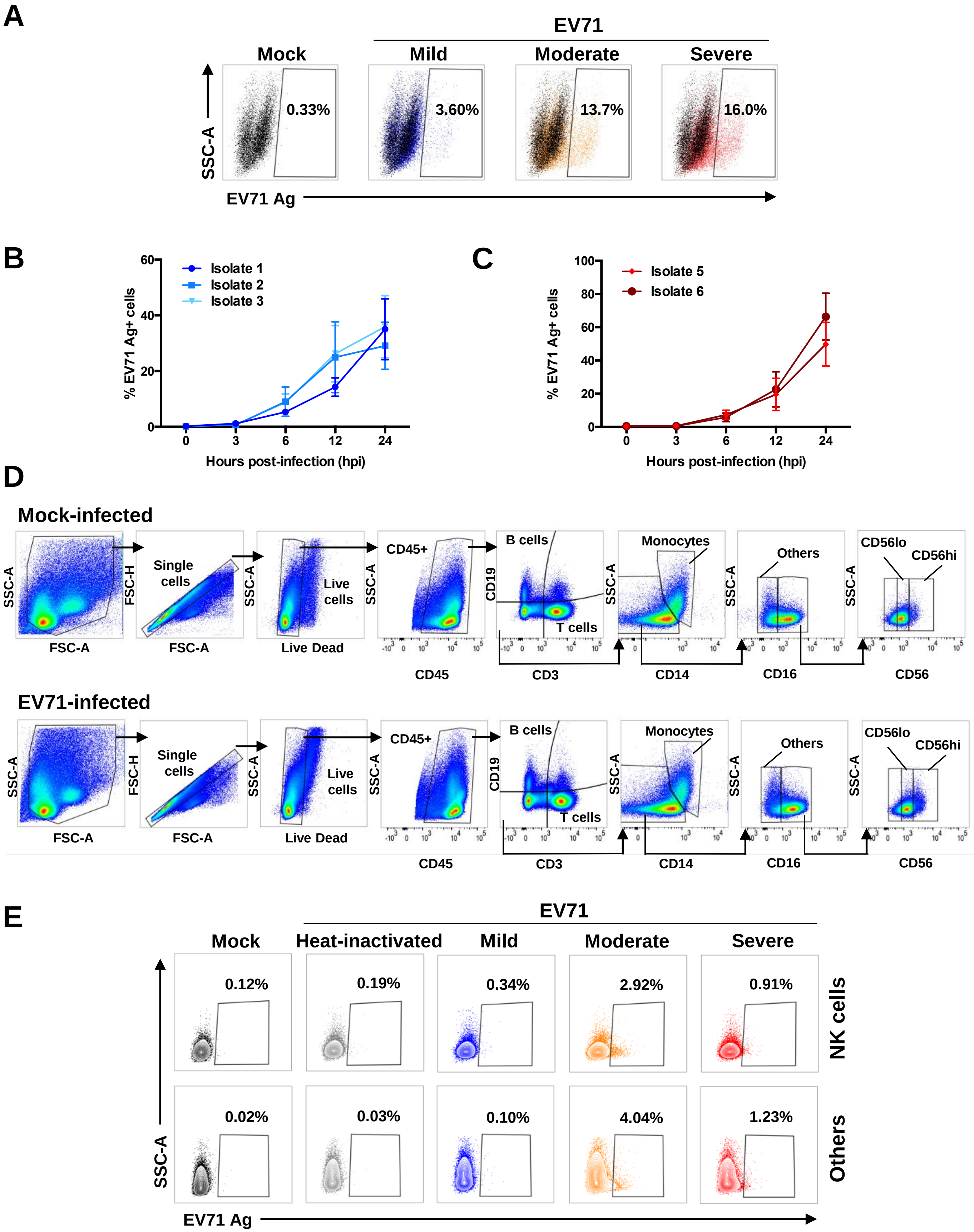


**Figure S2. EV71 infection in RD cells and primary human PBMCs.** (A) RD cells were infected with mild (isolate 1), moderate (isolate 4) and severe (isolate 5) EV71 at MOI 10 and harvested at 0, 3, 6, 12 and 24 hpi. Representative dot plots of EV71-infected RD cells at 12 hpi. (B-C) Quantification of VP1 antigen by flow cytometry of EV71 (B) isolates 1, 2 and 3, and (C) isolates 5 and 6 in RD cells at 0, 3, 6, 12 and 24 hpi at MOI 10. Data are presented as mean ± SEM and representative of 3 independent experiments. Statistical analysis was carried out with Kruskal-Wallis with Dunn's multiple comparisons test to compare among EV71 isolates at the respective time-points. (D-E) Human primary PBMCs were infected with mild, moderate, severe and heat-inactivated EV71 isolates at MOI 5 and harvested at 0, 6, 12 and 24 hpi. (D) Representative illustration of flow cytometry gating strategy from one donor at 6 hpi. (E) Representative contour plots of EV71-infected cell subsets at 12 hpi.

**Figure S2 Amrun *et al*., 2019**
