## Supplementary material for "TREM-1 activation is a key regulator in driving severe pathogenesis of enterovirus 71 infection": Figure S3

A

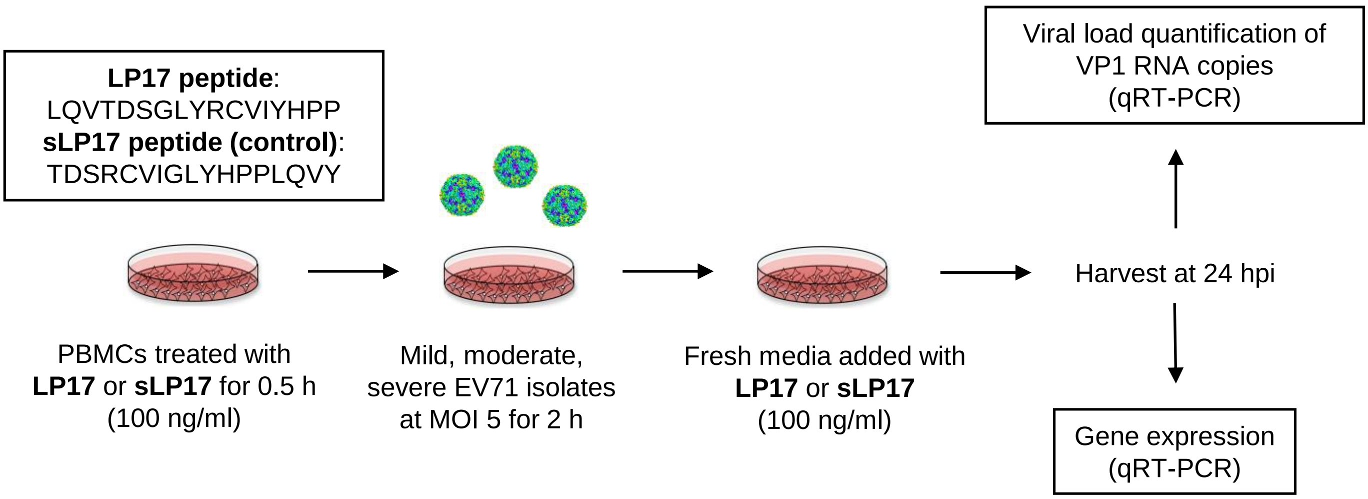


B

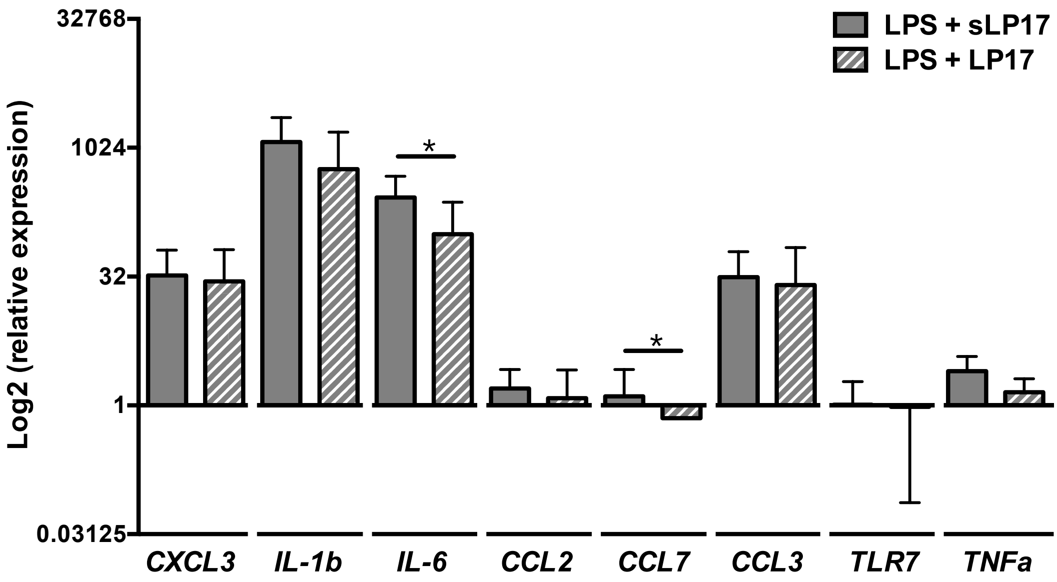


**Figure S3. Treatment of PBMCs with LP17 or sLP17 peptide.** Human primary PBMCs (n=7) were pre-treated with 100 ng/ml of LP17 or the control sLP17 peptide, before infection with mild, moderate, or severe EV71 isolates at MOI 5. Mock-infected and LPS-treated (50 ng/ml) PBMCs were used as negative and positive controls respectively. Cells were replenished with fresh media containing respective peptides and harvested at 24 hpi for gene expression and viral load quantifications. (A) Schematic illustration of the method. (B) Bar graphs showing the relative expression levels of genes involved in the TREM-1 pathway by qRT-PCR in the different peptide treatments of LPS-treated PBMCs. Data are presented as mean ± SD. Statistical analysis was carried out with Wilcoxon matched-pairs signed rank test (*$p$<0.05).

**Figure S3 Amrun *et al.*, 2019**
