## Supplementary material for "TREM-1 activation is a key regulator in driving severe pathogenesis of enterovirus 71 infection": Table S1

### Table S1: Percentage identity matrix of EV71 isolates

| Gene | Percentage identity (%)[a] | | | | | | |
|---|---|---|---|---|---|---|---|
| CDS | **Isolate** | **1** | **2** | **3** | **4** | **5** | **6** |
| | **1** | 100 | 99.41 | 96.44 | 96.41 | 96.46 | 96.41 |
| | **2** | 99.41 | 100 | 96.35 | 96.4 | 96.37 | 96.32 |
| | **3** | 96.44 | 96.35 | 100 | 96.55 | 99.98 | 99.64 |
| | **4** | 96.41 | 96.4 | 96.55 | 100 | 96.56 | 96.53 |
| | **5** | 96.46 | 96.37 | 99.98 | 96.56 | 100 | 99.65 |
| | **6** | 96.41 | 96.32 | 99.64 | 96.53 | 99.65 | 100 |
| VP4 | **Isolate** | **1** | **2** | **3** | **4** | **5** | **6** |
| | **1** | 100 | 100 | 96.14 | 95.65 | 96.14 | 95.65 |
| | **2** | 100 | 100 | 96.14 | 95.65 | 96.14 | 95.65 |
| | **3** | 96.14 | 96.14 | 100 | 96.14 | 100 | 99.52 |
| | **4** | 95.65 | 95.65 | 96.14 | 100 | 96.14 | 95.65 |
| | **5** | 96.14 | 96.14 | 100 | 96.14 | 100 | 99.52 |
| | **6** | 95.65 | 95.65 | 99.52 | 95.65 | 99.52 | 100 |
| VP2 | **Isolate** | **1** | **2** | **3** | **4** | **5** | **6** |
| | **1** | 100 | 98.95 | 95.67 | 97.11 | 95.67 | 96.19 |
| | **2** | 98.95 | 100 | 95.67 | 97.38 | 95.67 | 96.19 |
| | **3** | 95.67 | 95.67 | 100 | 96.19 | 100 | 99.48 |
| | **4** | 97.11 | 97.38 | 96.19 | 100 | 96.19 | 96.72 |
| | **5** | 95.67 | 95.67 | 100 | 96.19 | 100 | 99.48 |
| | **6** | 96.19 | 96.19 | 99.48 | 96.72 | 99.48 | 100 |
| VP3 | **Isolate** | **1** | **2** | **3** | **4** | **5** | **6** |
| | **1** | 100 | 98.9 | 96.42 | 96.42 | 96.42 | 96.28 |
| | **2** | 98.9 | 100 | 96.14 | 96.42 | 96.14 | 96.01 |
| | **3** | 96.42 | 96.14 | 100 | 96.83 | 100 | 99.59 |
| | **4** | 96.42 | 96.42 | 96.83 | 100 | 96.83 | 96.42 |
| | **5** | 96.42 | 96.14 | 100 | 96.83 | 100 | 99.59 |
| | **6** | 96.28 | 96.01 | 99.59 | 96.42 | 99.59 | 100 |
| VP1 | **Isolate** | **1** | **2** | **3** | **4** | **5** | **6** |
| | **1** | 100 | 99.66 | 96.97 | 96.75 | 96.97 | 97.31 |
| | **2** | 99.66 | 100 | 96.63 | 96.63 | 96.63 | 96.97 |
| | **3** | 96.97 | 96.63 | 100 | 96.18 | 100 | 99.66 |
| | **4** | 96.75 | 96.63 | 96.18 | 100 | 96.18 | 96.52 |
| | **5** | 96.97 | 96.63 | 100 | 96.18 | 100 | 99.66 |
| | **6** | 97.31 | 96.97 | 99.66 | 96.52 | 99.66 | 100 |
| 2A | **Isolate** | **1** | **2** | **3** | **4** | **5** | **6** |
| | **1** | 100 | 98.89 | 96.22 | 96.44 | 96.22 | 96.22 |
| | **2** | 98.89 | 100 | 95.78 | 95.78 | 95.78 | 95.78 |
| | **3** | 96.22 | 95.78 | 100 | 97.11 | 100 | 100 |
| | **4** | 96.44 | 95.78 | 97.11 | 100 | 97.11 | 97.11 |
| | **5** | 96.22 | 95.78 | 100 | 97.11 | 100 | 100 |
| | **6** | 96.22 | 95.78 | 100 | 97.11 | 100 | 100 |
| | **Isolate** | **1** | **2** | **3** | **4** | **5** | **6** |
| | **1** | 100 | 99.33 | 94.61 | 95.29 | 94.61 | 94.28 |

| | | | | | | | |
|---|---|---|---|---|---|---|---|
| **2B** | **2** | 99.33 | 100 | 95.29 | 95.96 | 95.29 | 94.95 |
| | **3** | 94.61 | 95.29 | 100 | 94.95 | 100 | 99.66 |
| | **4** | 95.29 | 95.96 | 94.95 | 100 | 94.95 | 94.61 |
| | **5** | 94.61 | 95.29 | 100 | 94.95 | 100 | 99.66 |
| | **6** | 94.28 | 94.95 | 99.66 | 94.61 | 99.66 | 100 |
| **2C** | **Isolate** | **1** | **2** | **3** | **4** | **5** | **6** |
| | **1** | 100 | 99.59 | 97.06 | 96.86 | 97.16 | 96.86 |
| | **2** | 99.59 | 100 | 97.06 | 96.86 | 97.16 | 96.86 |
| | **3** | 97.06 | 97.06 | 100 | 96.76 | 99.9 | 99.39 |
| | **4** | 96.86 | 96.86 | 96.76 | 100 | 96.86 | 96.66 |
| | **5** | 97.16 | 97.16 | 99.9 | 96.86 | 100 | 99.49 |
| | **6** | 96.86 | 96.86 | 99.39 | 96.66 | 99.49 | 100 |
| **3A** | **Isolate** | **1** | **2** | **3** | **4** | **5** | **6** |
| | **1** | 100 | 100 | 95.83 | 92.26 | 95.83 | 95.83 |
| | **2** | 100 | 100 | 95.83 | 92.26 | 95.83 | 95.83 |
| | **3** | 95.83 | 95.83 | 100 | 95.24 | 100 | 100 |
| | **4** | 92.26 | 92.26 | 95.24 | 100 | 95.24 | 95.24 |
| | **5** | 95.83 | 95.83 | 100 | 95.24 | 100 | 100 |
| | **6** | 95.83 | 95.83 | 100 | 95.24 | 100 | 100 |
| **3B** | **Isolate** | **1** | **2** | **3** | **4** | **5** | **6** |
| | **1** | 100 | 99.36 | 97.44 | 97.44 | 97.44 | 97.44 |
| | **2** | 99.36 | 100 | 98.08 | 98.08 | 98.08 | 98.08 |
| | **3** | 97.44 | 98.08 | 100 | 97.44 | 100 | 100 |
| | **4** | 97.44 | 98.08 | 97.44 | 100 | 97.44 | 97.44 |
| | **5** | 97.44 | 98.08 | 100 | 97.44 | 100 | 100 |
| | **6** | 97.44 | 98.08 | 100 | 97.44 | 100 | 100 |
| **3C** | **Isolate** | **1** | **2** | **3** | **4** | **5** | **6** |
| | **1** | 100 | 99.64 | 95.45 | 95.99 | 95.45 | 95.63 |
| | **2** | 99.64 | 100 | 95.45 | 95.99 | 95.45 | 95.63 |
| | **3** | 95.45 | 95.45 | 100 | 95.63 | 100 | 99.82 |
| | **4** | 95.99 | 95.99 | 97.63 | 100 | 97.63 | 97.81 |
| | **5** | 95.45 | 95.45 | 100 | 95.63 | 100 | 99.82 |
| | **6** | 95.63 | 95.63 | 99.82 | 97.81 | 99.82 | 100 |
| **3D** | **Isolate** | **1** | **2** | **3** | **4** | **5** | **6** |
| | **1** | 100 | 100 | 96.97 | 96.39 | 96.97 | 96.61 |
| | **2** | 99.57 | 100 | 96.83 | 96.25 | 96.83 | 96.46 |
| | **3** | 96.97 | 96.83 | 100 | 96.54 | 100 | 99.64 |
| | **4** | 96.39 | 96.25 | 96.54 | 100 | 96.54 | 96.32 |
| | **5** | 96.97 | 96.83 | 100 | 96.54 | 100 | 99.64 |
| | **6** | 100 | 96.46 | 99.64 | 96.32 | 99.64 | 100 |

[a]Percentage identity matrix of EV71 isolates based on the nucleotide sequences as determined by MUSCLE. Numbers highlighted in red are the lowest percentages in the respective regions.
