## Supplementary material for "TREM-1 activation is a key regulator in driving severe pathogenesis of enterovirus 71 infection": Table S2

**Table S2: List of common significant differentially expressed genes (DEGs) in EV71-infected PBMCs across time**

| Gene | Log$_2$-(fold change of infected over heat-inactivated) | | | | | | | | |
|---|---|---|---|---|---|---|---|---|---|
| | Mild EV71 | | | Moderate EV71 | | | Severe EV71 | | |
| | 6 hpi | 12 hpi | 24 hpi | 6 hpi | 12 hpi | 24 hpi | 6 hpi | 12 hpi | 24 hpi |
| *APOBEC3A* | 1.555 | 2.558 | 3.195 | 2.649 | 3.399 | 3.890 | 2.116 | 3.146 | 3.410 |
| *AXL* | 1.603 | 2.021 | 1.385 | 2.470 | 2.913 | 1.244 | 1.905 | 2.758 | 1.285 |
| *DDX58* | 1.754 | 1.576 | 1.860 | 2.167 | 2.193 | 1.934 | 1.873 | 1.912 | 1.855 |
| *DHX58* | 1.519 | 1.492 | 1.645 | 1.843 | 1.892 | 1.661 | 1.634 | 1.813 | 1.657 |
| *EIF2AK2* | 2.178 | 1.947 | 1.751 | 2.519 | 2.245 | 1.927 | 2.280 | 2.102 | 1.926 |
| *GMPR* | 1.770 | 2.384 | 2.602 | 2.281 | 3.035 | 2.309 | 1.635 | 2.826 | 2.224 |
| *HELZ2* | 1.853 | 1.835 | 2.036 | 2.164 | 2.201 | 2.311 | 1.862 | 2.255 | 2.300 |
| *HERC5* | 2.870 | 2.562 | 2.361 | 3.106 | 3.270 | 2.006 | 2.871 | 3.003 | 2.033 |
| *HERC6* | 2.489 | 2.089 | 2.358 | 2.724 | 2.624 | 2.512 | 2.450 | 2.447 | 2.534 |
| *IFI27* | 1.557 | 2.789 | 2.644 | 3.061 | 3.630 | 3.161 | 2.339 | 3.402 | 2.957 |
| *IFI35* | 1.175 | 1.385 | 1.530 | 1.601 | 1.792 | 1.479 | 1.286 | 1.556 | 1.351 |
| *IFI44* | 2.442 | 2.475 | 2.315 | 2.629 | 2.864 | 2.293 | 2.423 | 2.706 | 2.234 |
| *IFI44L* | 2.666 | 2.599 | 2.582 | 2.933 | 3.009 | 2.653 | 2.642 | 2.898 | 2.640 |
| *IFI6* | 2.632 | 2.779 | 2.314 | 2.963 | 3.286 | 2.355 | 2.431 | 2.922 | 2.094 |
| *IFIH1* | 1.281 | 1.425 | 1.560 | 1.766 | 1.864 | 1.619 | 1.534 | 1.706 | 1.546 |
| *IFIT5* | 1.634 | 1.462 | 1.445 | 1.869 | 1.976 | 1.487 | 1.761 | 1.796 | 1.512 |
| *IFITM1* | 1.632 | 1.420 | 2.207 | 1.983 | 2.145 | 2.317 | 1.937 | 2.019 | 2.317 |
| *IFITM3* | 1.424 | 2.174 | 1.849 | 2.199 | 2.787 | 1.876 | 1.767 | 2.538 | 1.755 |
| *IRF7* | 1.401 | 1.661 | 1.701 | 1.707 | 1.903 | 1.741 | 1.465 | 1.784 | 1.742 |
| *ISG15* | 2.654 | 3.075 | 3.472 | 3.411 | 3.928 | 3.134 | 2.978 | 3.573 | 3.058 |
| *ISG20* | 1.737 | 1.587 | 2.368 | 2.063 | 2.184 | 2.186 | 1.736 | 1.997 | 2.140 |
| *LY6E* | 1.903 | 1.992 | 1.801 | 2.209 | 2.434 | 1.884 | 1.915 | 2.178 | 1.747 |
| *MX1* | 3.061 | 2.640 | 2.265 | 3.517 | 3.297 | 2.326 | 3.207 | 3.106 | 2.242 |
| *MX2* | 2.870 | 2.419 | 2.145 | 3.380 | 2.938 | 2.321 | 3.129 | 2.840 | 2.248 |
| *NT5C3A* | 1.732 | 1.457 | 2.173 | 2.057 | 2.255 | 2.012 | 1.613 | 1.760 | 2.002 |
| *OASL* | 1.870 | 2.073 | 2.331 | 2.278 | 2.726 | 2.529 | 1.936 | 2.373 | 2.433 |
| *PLSCR1* | 1.786 | 1.727 | 1.828 | 2.152 | 2.167 | 1.849 | 1.906 | 1.980 | 1.746 |
| *PNPT1* | 1.943 | 1.573 | 1.718 | 2.183 | 2.285 | 1.578 | 2.079 | 2.027 | 1.600 |
| *RIN2* | 1.078 | 1.258 | 1.145 | 1.850 | 1.865 | 1.553 | 1.461 | 1.650 | 1.231 |
| *RSAD2* | 3.025 | 3.282 | 3.670 | 3.721 | 4.371 | 3.778 | 3.219 | 3.936 | 3.653 |
| *RTP4* | 1.378 | 1.415 | 1.707 | 1.488 | 2.072 | 1.395 | 1.266 | 1.651 | 1.485 |
| *SAMD9* | 1.718 | 1.476 | 1.563 | 1.972 | 2.005 | 1.699 | 1.744 | 1.802 | 1.673 |
| *SPATS2L* | 1.888 | 1.550 | 2.066 | 2.435 | 2.223 | 2.002 | 2.157 | 1.880 | 1.884 |
| *TNFSF10* | 0.978 | 1.508 | 1.918 | 1.596 | 2.152 | 1.822 | 1.176 | 2.043 | 1.862 |
| *USP18* | 3.564 | 3.171 | 2.863 | 4.084 | 4.094 | 3.009 | 3.755 | 3.742 | 2.857 |
| *XAF1* | 1.802 | 1.600 | 1.723 | 2.105 | 1.843 | 1.923 | 1.877 | 1.822 | 1.863 |
