## Supplementary material for "TREM-1 activation is a key regulator in driving severe pathogenesis of enterovirus 71 infection": Table S3

**Table S3: List of canonical pathways of common differentially expressed genes (DEGs) from EV71-infected PBMCs at 6-24 hpi**

| No. | Canonical Pathways | -log(*p* -value) | Ratio | Molecules |
|---|---|---|---|---|
| 1 | Interferon Signaling | 10.40 | 1.67E-01 | *IFITM3,MX1,IFI35,IFI6,IFITM1,ISG15* |
| 2 | Activation of IRF by Cytosolic Pattern Recognition Receptors | 7.09 | 8.20E-02 | *DHX58,IFIH1,IRF7,DDX58,ISG15* |
| 3 | Role of RIG-I-like Receptors in Antiviral Innate Immunity | 6.01 | 9.52E-02 | *DHX58,IFIH1,IRF7,DDX58* |
| 4 | Role of Pattern Recognition Receptors in Recognition of Bacteria and Viruses | 4.04 | 3.05E-02 | *IFIH1,IRF7,DDX58,EIF2AK2* |
| 5 | Salvage Pathways of Pyrimidine Ribonucleotides | 1.89 | 2.11E-02 | *EIF2AK2,APOBEC3A* |
| 6 | Salvage Pathways of Pyrimidine Deoxyribonucleotides | 1.84 | 1.25E-01 | *APOBEC3A* |
| 7 | Guanosine Nucleotides Degradation III | 1.67 | 8.33E-02 | *NT5C3A* |
| 8 | Urate Biosynthesis/Inosine 5'-phosphate Degradation | 1.63 | 7.69E-02 | *NT5C3A* |
| 9 | Adenosine Nucleotides Degradation II | 1.57 | 6.67E-02 | *NT5C3A* |
| 10 | Purine Nucleotides Degradation II (Aerobic) | 1.49 | 5.56E-02 | *NT5C3A* |
| 11 | Role of Lipids/Lipid Rafts in the Pathogenesis of Influenza | 1.45 | 5.00E-02 | *RSAD2* |
| 12 | NAD Salvage Pathway II | 1.43 | 4.76E-02 | *NT5C3A* |
