## Supplementary material for "TREM-1 activation is a key regulator in driving severe pathogenesis of enterovirus 71 infection": Table S4

**Table S4: List of significant differentially expressed genes (DEGs) in EV71-infected PBMCs at 24 hpi**

| Gene | $Log_2$-(fold change of infected over heat-inactivated) | | |
|---|---|---|---|
| | Mild EV71 | Moderate EV71 | Severe EV71 |
| *AGRN* | 1.194 | 1.330 | 1.160 |
| *APOBEC3A* | 3.195 | 3.890 | 3.410 |
| *APOE* | -2.167 | -3.071 | -4.181 |
| *ATF3* | 1.251 | 1.413 | 1.276 |
| *AXL* | 1.385 | 1.244 | 1.285 |
| *B3GNT2* | 1.134 | 1.274 | 1.060 |
| *CCL3* | 1.319 | 1.582 | 1.640 |
| *CCL7* | 2.739 | 2.770 | 2.871 |
| *CCL8* | 3.567 | 4.017 | 3.842 |
| *CCR2* | -1.588 | -2.348 | -1.684 |
| *CFAP43* | 4.377 | 4.627 | 4.269 |
| *CLEC10A* | -1.242 | -1.862 | -1.137 |
| *CTSL* | 1.696 | 2.066 | 1.719 |
| *CXCL10* | 3.666 | 3.480 | 3.405 |
| *CYP19A1* | 5.404 | 6.382 | 5.805 |
| *DDX58* | 1.860 | 1.934 | 1.855 |
| *DDX60L* | 1.786 | 1.775 | 1.754 |
| *DHX58* | 1.645 | 1.661 | 1.657 |
| *DUSP5* | 0.998 | 1.218 | 1.309 |
| *EIF2AK2* | 1.751 | 1.927 | 1.926 |
| *EPSTI1* | 1.373 | 1.407 | 1.325 |
| *ERICH3* | 4.493 | 4.872 | 4.839 |
| *GMPR* | 2.602 | 2.309 | 2.224 |
| *HELZ2* | 2.036 | 2.311 | 2.300 |
| *HERC5* | 2.361 | 2.006 | 2.033 |
| *HERC6* | 2.358 | 2.512 | 2.534 |
| *HESX1* | 4.170 | 4.071 | 3.718 |
| *HPSE* | 1.750 | 1.859 | 1.633 |
| *IFI27* | 2.644 | 3.161 | 2.957 |
| *IFI35* | 1.530 | 1.479 | 1.351 |
| *IFI44* | 2.315 | 2.293 | 2.234 |
| *IFI44L* | 2.582 | 2.653 | 2.640 |
| *IFI6* | 2.314 | 2.355 | 2.094 |

| | | | |
|---|---|---|---|
| *IFIH1* | 1.560 | 1.619 | 1.546 |
| *IFIT5* | 1.445 | 1.487 | 1.512 |
| *IFITM1* | 2.207 | 2.317 | 2.317 |
| *IFITM3* | 1.849 | 1.876 | 1.755 |
| *IGFBP4* | 1.421 | 1.164 | 1.223 |
| *IL1RN* | 2.456 | 2.534 | 2.299 |
| *IL27* | 2.644 | 2.521 | 2.321 |
| *IRF7* | 1.701 | 1.741 | 1.742 |
| *ISG15* | 3.472 | 3.134 | 3.058 |
| *ISG20* | 2.368 | 2.186 | 2.140 |
| *KITLG* | 1.836 | 2.548 | 1.961 |
| *LAG3* | 1.740 | 1.569 | 1.849 |
| *LAMP3* | 1.114 | 1.260 | 1.506 |
| *LGALS3BP* | 1.127 | 1.213 | 1.304 |
| *LILRA5* | 1.510 | 1.794 | 1.746 |
| *LY6E* | 1.801 | 1.884 | 1.747 |
| *MX1* | 2.265 | 2.326 | 2.242 |
| *MX2* | 2.145 | 2.321 | 2.248 |
| *NEURL3* | 2.961 | 3.867 | 3.692 |
| *NEXN* | 2.611 | 2.154 | 2.004 |
| *NT5C3A* | 2.173 | 2.012 | 2.002 |
| *OASL* | 2.331 | 2.529 | 2.433 |
| *OTOF* | 2.797 | 3.475 | 3.244 |
| *PADI2* | -1.964 | -2.652 | -2.116 |
| *PHF11* | 1.047 | 2.025 | 2.233 |
| *PLSCR1* | 1.828 | 1.849 | 1.746 |
| *PNPT1* | 1.718 | 1.578 | 1.600 |
| *RIN2* | 1.145 | 1.553 | 1.231 |
| *RSAD2* | 3.670 | 3.778 | 3.653 |
| *RTP4* | 1.707 | 1.395 | 1.485 |
| *SAMD4A* | 1.198 | 1.340 | 1.254 |
| *SAMD9* | 1.563 | 1.699 | 1.673 |
| *SAMD9L* | 1.539 | 1.631 | 1.526 |
| *SLC38A5* | 1.587 | 1.798 | 1.754 |
| *SPATS2L* | 2.066 | 2.002 | 1.884 |
| *TNFSF10* | 1.918 | 1.822 | 1.862 |
| *USP18* | 2.863 | 3.009 | 2.857 |
| *XAF1* | 1.723 | 1.923 | 1.863 |
| *ZBP1* | 1.459 | 1.541 | 1.535 |

| | | | |
|---|---|---|---|
| *CHI3L1* | -1.672 | -2.382 | N.A. |
| *EPHB2* | 0.984 | 1.012 | N.A. |
| *HSPA1A* | 1.287 | 1.001 | N.A. |
| *IL1R2* | 1.574 | 1.109 | N.A. |
| *RASGEF1B* | 1.127 | 1.194 | N.A. |
| *ABTB2* | N.A. | 1.930 | 1.891 |
| *ALDH1A1* | N.A. | -1.559 | -1.741 |
| *ALDH2* | N.A. | -1.291 | -1.430 |
| *ANKRD1* | N.A. | 3.335 | 3.245 |
| *ARNT2* | N.A. | 2.335 | 2.118 |
| *C19orf66* | N.A. | 1.094 | 1.145 |
| *CCL4* | N.A. | 1.241 | 1.539 |
| *CD300E* | N.A. | 2.072 | 1.738 |
| *CD9* | N.A. | -1.004 | -1.546 |
| *CLMP* | N.A. | 3.837 | 3.592 |
| *FBP1* | N.A. | -1.194 | -1.283 |
| *FSCN1* | N.A. | 1.174 | 1.103 |
| *IRG1* | N.A. | 2.175 | 1.995 |
| *ITGB8* | N.A. | 1.932 | 2.388 |
| *LTA4H* | N.A. | -1.325 | -1.423 |
| *MDK* | N.A. | 2.054 | 2.072 |
| *NFAM1* | N.A. | -1.043 | -1.037 |
| *PLBD1* | N.A. | -1.145 | -1.165 |
| *SERPINA1* | N.A. | -1.193 | -1.228 |
| *SERPINB2* | N.A. | 1.169 | 1.735 |
| *SSTR3* | N.A. | 1.631 | 1.831 |
| *TNFAIP6* | N.A. | 1.611 | 1.699 |
| *TREM2* | N.A. | -1.444 | -2.012 |
| *WASH5P* | N.A. | 5.643 | 5.554 |
| *OLR1* | -1.552 | N.A. | -1.763 |
| *DYNLT1* | 1.195 | N.A. | N.A. |
| *GCH1* | 1.016 | N.A. | N.A. |
| *HSPA1B* | 1.109 | N.A. | N.A. |
| *MARCKS* | 1.186 | N.A. | N.A. |
| *MT2A* | 1.392 | N.A. | N.A. |
| *NMI* | 0.964 | N.A. | N.A. |
| *NUPR1* | 2.771 | N.A. | N.A. |
| *PGAP1* | 2.100 | N.A. | N.A. |
| *PPBP* | 1.347 | N.A. | N.A. |

| | | | |
|---|---|---|---|
| *RNASE1* | 1.655 | N.A. | N.A. |
| *RTCB* | 1.038 | N.A. | N.A. |
| *SCIN* | 1.662 | N.A. | N.A. |
| *SPP1* | 1.253 | N.A. | N.A. |
| *ATP13A2* | N.A. | 1.481 | N.A. |
| *CCL24* | N.A. | -1.378 | N.A. |
| *CD1D* | N.A. | -1.425 | N.A. |
| *CXCL13* | N.A. | 2.369 | N.A. |
| *FCN1* | N.A. | -1.242 | N.A. |
| *IDO1* | N.A. | 1.318 | N.A. |
| *MSR1* | N.A. | 1.128 | N.A. |
| *MYCL* | N.A. | -1.278 | N.A. |
| *MYO10* | N.A. | 1.705 | N.A. |
| *PML* | N.A. | 1.156 | N.A. |
| *RNF19B* | N.A. | 1.097 | N.A. |
| *SCARB1* | N.A. | -1.575 | N.A. |
| *SEMA4A* | N.A. | 0.997 | N.A. |
| *SLC1A4* | N.A. | 1.013 | N.A. |
| *SLC39A8* | N.A. | 1.525 | N.A. |
| *SLC7A8* | N.A. | 1.431 | N.A. |
| *SPARC* | N.A. | -2.077 | N.A. |
| *SRGAP2* | N.A. | 1.143 | N.A. |
| *TIFAB* | N.A. | -1.290 | N.A. |
| *XCR1* | N.A. | 1.485 | N.A. |
| *ABCC3* | N.A. | N.A. | -1.363 |
| *APOL4* | N.A. | N.A. | -1.196 |
| *CCL4L1* | N.A. | N.A. | 1.636 |
| *CEBPA* | N.A. | N.A. | -1.142 |
| *CKB* | N.A. | N.A. | 2.148 |
| *CXCL1* | N.A. | N.A. | 1.304 |
| *CXCL3* | N.A. | N.A. | 1.036 |
| *CXCL5* | N.A. | N.A. | 1.526 |
| *FN1* | N.A. | N.A. | -1.274 |
| *IER5L* | N.A. | N.A. | -1.502 |
| *IL-1β* | N.A. | N.A. | 1.083 |
| *IL-6* | N.A. | N.A. | 3.212 |
| *PHF11* | N.A. | N.A. | 1.012 |
| *PRAM1* | N.A. | N.A. | -1.526 |
| *PYCR1* | N.A. | N.A. | 2.920 |

| | | | |
|---|---|---|---|
| *RGS1* | N.A. | N.A. | 1.075 |
