## Supplementary material for "TREM-1 activation is a key regulator in driving severe pathogenesis of enterovirus 71 infection": Table S5

**Table S5: List of canonical pathways from differentially expressed genes (DEGs) of PBMC-infected EV71 isolates at 24 hpi**

| No. | Canonical Pathways | Activation *z* -score | | | -log(*p* -value) | | | Molecules |
|---|---|---|---|---|---|---|---|---|
| | | Mild EV71 | Moderate EV71 | Severe EV71 | Mild EV71 | Moderate EV71 | Severe EV71 | |
| 1 | Interferon Signaling | 2.449 | 2.449 | 2.449 | 7.894 | 7.143 | 7.322 | *IFITM3, MX1, IFI6, IFI35, IFITM1, ISG15* |
| 2 | LXR/RXR Activation | -2.000 | -2.236 | -2.449 | 2.651 | 4.029 | 4.196 | *APOE, IL1RN, IL-1b, SERPINA1, IL-6, CCL7* |
| 3 | Role of Pattern Recognition Receptors in Recognition of Bacteria and Viruses | 2.000 | 2.000 | 2.449 | 2.526 | 2.084 | 4.004 | *IFIH1, IRF7, DDX58, IL-1b, IL-6, EIF2AK2* |
| 4 | IL-6 Signaling | N.A. | 2.000 | 2.236 | 1.638 | 2.050 | 3.010 | *TNFAIP6, IL1RN, CYP19A1, IL1-b, IL-6* |
| 5 | Role of RIG-I-like Receptors in Antiviral Innate Immunity | 1.000 | 1.000 | 1.000 | 4.413 | 3.925 | 4.042 | *IFIH1, DHX58, IRF7, DDX58* |
| 6 | Activation of IRF by Cytosolic Pattern Recognition Receptors | 0.816 | 0.816 | 1.134 | 6.479 | 5.741 | 7.293 | *IFIH1, DHX58, IRF7, ZBP1, DDX58, IL-6, ISG15* |
| 7 | Neuro-inflammation Signaling Pathway | N.A. | 1.000 | 1.633 | 0.806 | 0.960 | 2.142 | *CXCL10, IRF7, TREM2, IL-1b, IL-6, CCL3* |
| 8 | Role of IL-17F in Allergic Inflammatory Airway Diseases | N.A. | N.A. | 2.449 | 1.809 | 2.682 | 8.464 | *CXCL10, CCL4, IL-1b, CXCL1, CXCL5, IL-6, CCL7* |
| 9 | TREM-1 Signaling | N.A. | N.A. | 2.236 | 1.396 | 1.174 | 4.327 | *CXCL3, IL-1b, IL6, CCL3, CCL7* |
| 10 | Dendritic Cell Maturation | N.A. | 0.000 | 1.342 | 0.244 | 1.586 | 2.399 | *IL1RN, FSCN1, TREM2, IL-1b, IL-6* |
