## Supplementary material for "TREM-1 activation is a key regulator in driving severe pathogenesis of enterovirus 71 infection": Table S6

**Table S6: Information of primers used for gene expression**

| Gene | Direction | Sequence (5' to 3') |
|---|---|---|
| *CXCL3* | Forward | CGCCCAAACCGAAGTCATAG |
| | Reverse | GCTCCCCTTGTTCAGTATCTTTT |
| *IL-1β* | Forward | AAATACCTGTGGCCTTGGGC |
| | Reverse | TTTGGGATCTACACTCTCCAGCT |
| *IL-6* | Forward | CACAGACAGCCACTCACCTCTTCAGAACGA |
| | Reverse | ACCAGTGATTTTCACCAGGCAAGTCTC |
| *CCL2* | Forward | CAGCCAGATGCAATCAATGCC |
| | Reverse | TGGAATCCTGAACCCACTTCT |
| *CCL7* | Forward | TGCTCAGCCAGTTGGGATTA |
| | Reverse | GGACAGTGGCTACTGGTGGT |
| *CCL3* | Forward | TCAGACTTCAGAAGGACACGG |
| | Reverse | CTGCATGATTCTGAGCAGGTG |
| *TLR7* | Forward | TCCTTGGGGCTAGATGGTTTC |
| | Reverse | TCCACGATCACATGGTTCTTTG |
| *TNFα* | Forward | TGCTTGTTCCTCAGCCTCTT |
| | Reverse | GGAAGACCCCTCCCAGATAG |
| *GAPDH* | Forward | CCACATCGCTCAGACACCAT |
| | Reverse | GGCAACAATATCCACTTTACCAGAGT |
